# DeCoDe: degenerate codon design for complete protein-coding DNA libraries

**DOI:** 10.1101/809004

**Authors:** Tyler C. Shimko, Polly M. Fordyce, Yaron Orenstein

## Abstract

**Motivation:** High-throughput protein screening is a critical technique for dissecting and designing protein function. Libraries for these assays can be created through a number of means, including targeted or random mutagenesis of a template protein sequence or direct DNA synthesis. However, mutagenic library construction methods often yield vastly more non-functional than functional variants and, despite advances in large-scale DNA synthesis, individual synthesis of each desired DNA template is often prohibitively expensive. Consequently, many protein screening libraries rely on the use of degenerate codons (DCs), mixtures of DNA bases incorporated at specific positions during DNA synthesis, to generate highly diverse protein variant pools from only a few low-cost synthesis reactions. However, selecting DCs for sets of sequences that covary at multiple positions dramatically increases the difficulty of designing a DC library and leads to the creation of many undesired variants that can quickly outstrip screening capacity.

**Results:** We introduce a novel algorithm for total DC library optimization, DeCoDe, based on integer linear programming. DeCoDe significantly outperforms state-of-the-art DC optimization algorithms and scales well to more than a hundred proteins sharing complex patterns of covariation (*e.g.* the lab-derived avGFP lineage). Moreover, DeCoDe is, to our knowledge, the first DC design algorithm with the capability to encode mixed-length protein libraries. We anticipate DeCoDe to be broadly useful for a variety of library generation problems, ranging from protein engineering attempts that leverage mutual information to the reconstruction of ancestral protein states.

**Availability:** github.com/OrensteinLab/DeCoDe

## Introduction

A protein’s function is inextricably linked to its amino acid sequence. Stability, flexibility, enzymatic turnover, and binding affinity all depend directly on the specific structures available to the sequence of a given protein (9, 25). Due to the critical importance of protein products in medicine and industry, a number of high-throughput screening techniques, including cell-surface display (3, 5, 10, 32), phage display (37)), mRNA display (31, 40), and droplet-based enzyme screens (1, 33) have been developed to link protein sequence to function. The majority of these techniques allow directed evolution for the development of functional properties. These techniques all require starting libraries of DNA encoding a vast number of protein variants, generally > 10^6^. Consequently, a number of strategies have been developed to enrich screening libraries for protein variants with a high likelihood of functionality.

Historically, targeted protein library design has relied on the expertise of trained biochemists. However, beginning in the early 2000s, computational methods for linking protein sequence to function started to emerge. Initial methods employed biophysical models of protein structure and enabled computational modeling of hundreds to thousands of protein variants prior to experimental screening (16, 19). More recent developments employ machine learning to rapidly predict the function of a novel protein sequence without the computational expense of simulating protein dynamics. These machine learning methods predict function for large, targeted protein libraries of a size on par with the throughput of screening methods (7, 34, 45). In many cases, these libraries encode covarying sequence structures to maximize the number of properly folded functional variants (15, 21). Therefore, testing these predictions in experimental screens requires the ability to quickly and cheaply generate pools of DNA templates with covarying residues that code for protein variants.

The gold standard for DNA library construction is direct DNA synthesis of each individual library member followed by pooling (4). This strategy guarantees the inclusion of each desired DNA sequence without introducing off-target constructs. However, direct synthesis is often prohibitively expensive for large libraries and has limitations on both the length of each synthesized construct and the total number of constructs that can be synthesized in parallel. While recent advances in DNA synthesis technology are enabling direct and specific synthesis for longer and larger libraries (20, 26, 29), the number of possible full length protein-coding constructs (10^3^ - 10^6^) remains several orders of magnitude below the desired library sizes for most protein screening methods (10^6^ - 10^13^).

Methods that exploit the redundancy of the genetic code to generate large, semi-targeted libraries balance low cost, simple production with library output sizes suitable for protein screening. A degenerate codon (DC) is a mixture of nucleotide triplets capable of collectively encoding more than one amino acid (2, 17, 44). DC libraries combine mixtures of nucleotides at specific positions during DNA synthesis to ultimately allow expression of protein mixtures from a single pooled DNA synthesis reaction. The composition of the nucleotide mixture incorporated at a given position allows DCs to be designed such that they include only specific subsets of the codon table, giving experimenters tighter control over the identity of the protein mixture than with simple random library construction. However, as the identity of the codon generated within each DNA construct is independent of every other DC, including a large number of DCs can quickly generate a library of possible DNA and protein sequences too large to be screened.

A number of groups have developed computational methods to maximize the number of desired target sequences created using DCs under a user-specified library size. Unfortunately, this problem is NP-hard (27, 28), meaning that a fast (polynomial time) algorithmic solution is extremely unlikely to exist. Instead, researchers have had to rely on a variety of heuristics or relaxations of the problem to develop algorithms that efficiently design DC libraries. One of the first methods to allow simultaneous optimization of all covarying sites was LibDesign (24). LibDesign optimizes DC usage to include as many complete targeted protein sequences as possible in the final library under a limit on the total number of sequences produced. As a result, libraries designed by LibDesign directly account for the three main structures of protein covariation as shown in Figures 1A, 1B, and 1C. However, because this algorithm relies on brute-force search, it is computationally inefficient and intractable for modern protein library designs that may vary at dozens or more positions. Moreover, LibDesign is incapable of accounting for length variation within the target library or using multiple synthesis reactions (i.e. a DNA library comprised of multiple sublibraries with each sublibrary produced according to a separate DNA template) to cover more targets without incurring large library size penalties.

**Fig 1.**
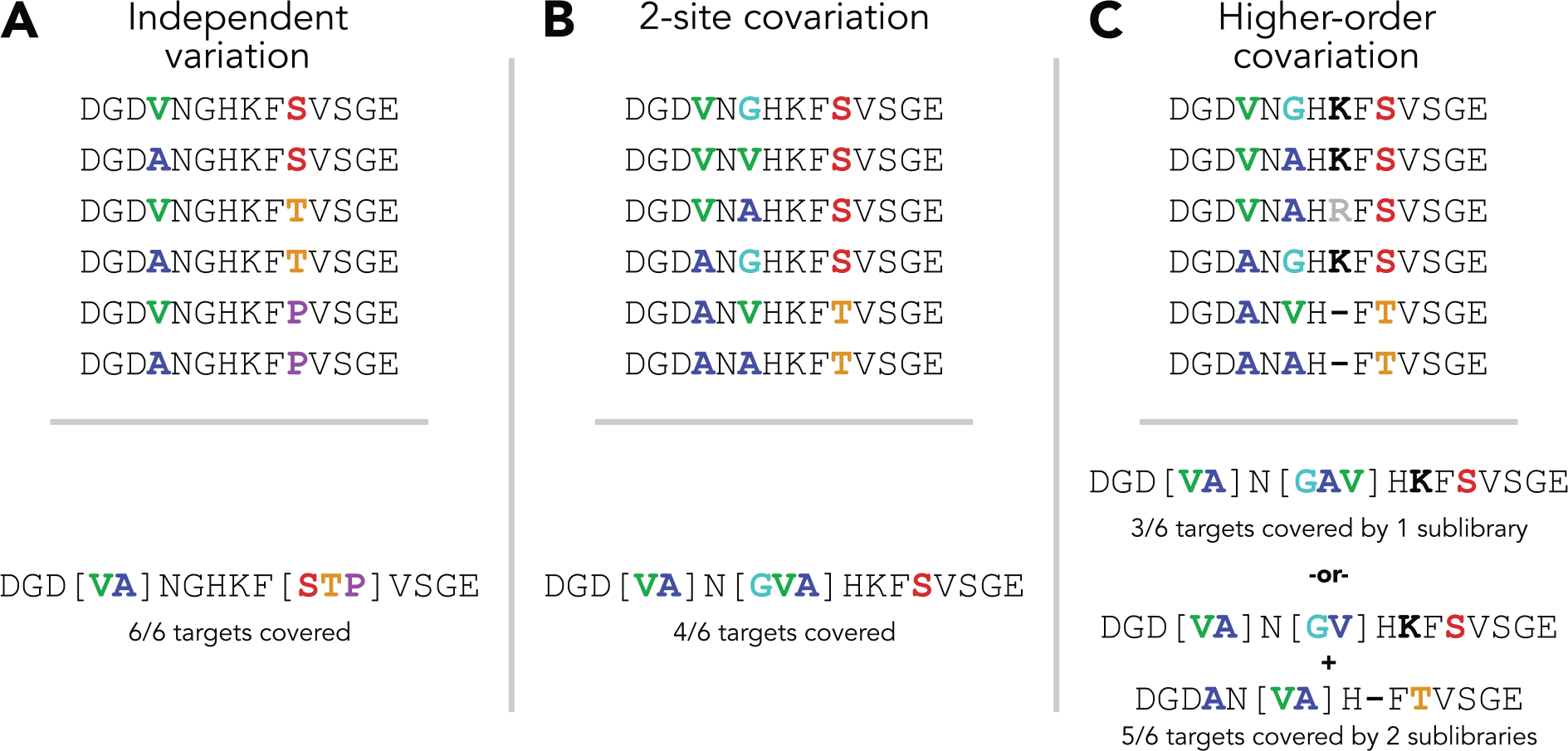
Examples of covariation structures within protein libraries (top row). For each covariation pattern, an optimal library was generated with a total size limit of 6 DNA sequences (bottom row). The amino acid sequences encoded by this library, along with the total number of targets covered is shown. **(A)** A library with two independently varying positions is shown (top). The DC library generated for this set of sequences was able to code all 6 targets under the total library size limit of 6. **(B)** A library with two covarying positions is shown (top). The DC library generated for this set of sequences was able to code only 4 targets under the total library size limit of 6. Note that the third variable position is not degenerate in the optimal library. **(C)** A library with multiple covarying positions and an indel mutation is shown (top). The DC library generated for this set of sequences was able to code only 3 targets under the total library size limit of 6. By adding a second sublibrary, the indel mutation could be handled by templates with different lengths and the total coverage increased to 5 targets.

As an alternative to total-library optimization, Optimization of Combinatorial Mutagenesis (OCoM) measures library quality by the maintenance of single and pairwise mutational frequencies (Figures 1A and 1B) and optimizes libraries using integer linear programming (ILP) (27). However, the pairwise sequence potential employed by OCoM fails to capture higherorder covariation patterns (Figure 1C) that would be explicitly considered under LibDesign’s objective. Such patterns are particularly important for functional networks within proteins (13, 23, 38) and could prove critical for the generation of libraries optimized for inclusion of functional variants. Furthermore, OCoM cannot design libraries of multiple lengths.

Due to the problem’s hardness, some groups have introduced relaxations to solve it in polynomial time with respect to the input size. SwiftLib is a DC optimization algorithm based on dynamic programming (DP) (14). To reduce computational complexity, SwiftLib considers each position of the protein library independently. Critically, SwiftLib can include multiple DCs at a given position to better cover the target library while staying under the same diversity limit. This strategy dramatically reduces the total size of the final library by minimizing or eliminating off-target amino acid inclusion at a given position. Despite these advantages, the simplification of the DC design problem used by SwiftLib eliminates its ability to account for covariation. While SwiftLib allows for the use of multiple degenerate DNA templates, it does not account for a gap and, therefore, lacks the ability to encode mixed length libraries.

Here, we present Degenerate Codon Design (DeCoDe), an algorithm for total-library DC optimization based on ILP that simultaneously addresses three critical gaps in current algorithmic solutions to DC library design: (i) direct accounting for high-order covariation, (ii) optimization over mixed length libraries, and (iii) inclusion of multiple degenerate DNA templates to cover more proteins of the input library under experimental constraints. For small libraries, DeCoDe often achieves optimal library designs in a reasonable amount of time (hours to days). For larger libraries, DeCoDe will output a feasible solution after a given time limit, and will report the gap between the current best number of targets covered and an upper bound on the optimal design. Because of its distinct advantages, we expect DeCoDe to be applicable to numerous protein engineering challenges including library design for directed evolution, reconstruction of ancestral protein states, and high-throughput biochemical analysis.

## Methods

### Preliminaries

We start with a few formal definitions that will aid in the problem definition and ILP formulation. Proofs supporting all ILP con-straints as well as the complete ILP formulations can be found in the Supplementary Information.

#### Definition 1

An **amino acid sequence** is a string over the amino acid alphabet Σ_*aa*_ plus a gap character indicating the absence of an amino acid at a given position.

#### Definition 2

A **codon** is a DNA triplet coding a single amino acid or stop or a gap character indicating the absence of a DNA triplet at a given position.

#### Definition 3

A **degenerate nucleotide** represents a subset of {*A*, *C*, *G*, *T*}.

In general, a symbol in an alphabet is said to be *degenerate* if it represents a set of symbols within the same alphabet and that set has a cardinality greater than one. Nucleotide degeneracy allows DNA molecules to be synthesized with a mixture of nucleotides at one or more specified positions, giving direct rise to degenerate codons.

#### Definition 4

A **degenerate codon** is a triple of degenerate nucleotides and, thus, codes a subset of Σ_*aa*_.

Whereas non-degenerate codons code one and only one amino acid residue or stop, degenerate codons can code multiple amino acids or stops by representing a mixture of non-degenerate codons. If we denote the DNA triplets represented by degenerate codon *x* as span_DNA_(*x*), then *x covers* non-degenerate codon *c* if and only if *c* ∈ span_DNA_(*x*).

#### Definition 5

A degenerate codon **covers** the DNA triplets represented by it and codes the amino acids they code.

We denote the amino acids encoded by degenerate codon *x* by span_AA_(*x*), then *x* codes *a* if and only if *a* ∈ span_AA_(*x*).

#### Definition 6

Definition 6: A sequence of degenerate codons, a degenerate template *G* = {*g*_1_, …, *g*_*P*_}, **codes** a sequence of amino acids *A* = {*a*_1_, …, *a*_*P*_} if and only if *a*_*p*_ ∈ span_AA_(*g*_*p*_) ∀1 ≤ *p* ≤ *P*.

A sequence of degenerate codons can be assembled to cover a DNA library coding a large number of protein sequences. We denote by span_*AA*_(*G*) the set of amino acid sequences coded by degenerate template *G*.

#### Definition 7

span_DNA_(*G*) of degenerate template *G* is called a **sublibrary**.

We note that the cardinality of span_DNA_(*G*) will grow exponentially with the number of degenerate codons included in *G*, quickly outstripping the capacity of all available experimental screening methods. Therefore, it is necessary to constrain the maximum number of non-degenerate DNA sequences produced, i.e. limit the cardinality of span_DNA_(*G*).

#### Definition 8

**A library** is made up of one or more sublibraries, each covered by a single degenerate template *G*_*s*_. Formally, a library is denoted by ∪_*s*_span_DNA_(*G*_*s*_).

By grouping similar sequences together in a sublibrary, each covered by a single degenerate template, then combining the sublibraries to generate the final library, more protein sequences can be encoded under the same DNA diversity limit.

### Problem definition

We define our problem as that of taking a set of desired protein sequences and producing a degenerate template coding as many of those sequences as possible while limiting the total number of DNA sequences covered as shown in Figures 1A and 1B.

### Max-coverage degenerate single-template design (MC-DSTD)

- INSTANCE: Set *T* of *P*-long amino acid sequences {*T*_*i*_} and DNA library size limit *M*.
- VALID SOLUTION: *P*-long degenerate template *G* s.t. |span_DNA_(*G*)| ≤ *M*.
- GOAL: Maximize |span_AA_(*G*) ∩*T*|

Optionally, we allow the library to be constructed as a combination of smaller sublibraries (Figure 1C), each covered by a single degenerate template, and coding a fraction of the total number of targeted proteins.

### Max-coverage degenerate multi-template design (MC-DMTD)

- INSTANCE: Set *T* of amino acid sequences {*T*_*i*_} with a maximum length of *P*, DNA library size limit *M*, and number of degenerate templates *g*.
- VALID SOLUTION: Set *G* of *P*-long degenerate templates {*G*_*s*_}, where |*G*| = *g*, such that 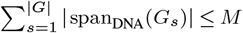.
- GOAL: Maximize 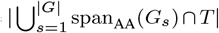.

Proteins of multiple lengths are allowed only in the multiple template design, as a single degenerate template codes for proteins of only one specific length. By using *x* gap codons in a template of length *P*, a template codes proteins of length *P*−*x*.

As noted by Parker and colleagues (27), it follows from the NP-hardness of the protein design problem (28) that the design of degenerate sequences varying non-independently at multiple positions is NP-hard. This finding holds whether designing a single template or multiple degenerate templates. Consequently, we devise a solution to this problem using integer linear programming (ILP), a method with efficient solvers that was applied to a myriad of NP-hard problems.

### MC-DSTD: Single library formulation

We first present a solution to the problem in which we cover the library using only a single sublibrary (one degenerate template). This restriction simplifies the calculation of the total produced library size |span_DNA_(*G*)|, which ordinarily requires multiplication of independent variables, an operation that is disallowed in linear programs.

#### Objective

To address this problem, we introduce the objective function:

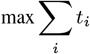

where *i* indexes an indicator variable such that *t*_*i*_ denotes whether target protein sequence *T*_*i*_ in the target library *T* can be translated from the set of DNA sequences *G* created by the protein sequence library. Formally, *t*_*i*_ = 1 ⇔ *T*_*i*_ ∈ span_AA_(*G*). By optimizing for inclusion of full length sequences, DeCoDe implicitly captures covariation structures within the library (Figures 1B and 1C).

### Single degenerate codon per position constraint

We introduce the variable *G*_*spd*_ where *s* denotes the index of the degenerate template, *p* denotes the position of the degenerate codon in the degenerate template with *P* representing the total number of positions, and *d* denotes the use of the *d*th degenerate codon at that specified position. Note that here, the MCDSTD problem, *s* = 1 as we are only considering the special case of having a single degenerate template cover the entire library. Upon the variable *G*_*spd*_, we introduce the following constraint:

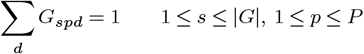

so that only a single degenerate codon can be employed at each position of each degenerate template.

#### Coverage constraints

We introduce integer matrix *D* and binary matrix 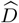. In matrix *D*, *D*_*da*_ corresponds to the number of non-degenerate codons covered by degenerate codon *d* that code the *a*th amino acid, gap, or stop. Matrix 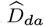 is a binary copy of matrix *D* where each value 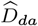 is the evaluated truth of *D*_*da*_ > 0. Variable *C*_*spa*_ is an indicator variable for coding the *a*th amino acid at position *p* by degenerate template *G*_*s*_. Therefore, the following relationship exists between *G* and *C* variables:

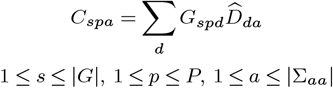

To indicate whether a specific target sequence *T*_*i*_ can be encoded by the sublibrary set, we introduce the variable *X*_*is*_. *X*_*is*_ indicates whether degenerate template *G*_*s*_ can code the target protein sequence indexed by *i*. We introduce the following constraints upon this variable, where *O* is defined such that *O*_*ipa*_ is a one-hot encoded representation of the target sequence *T*_*i*_:

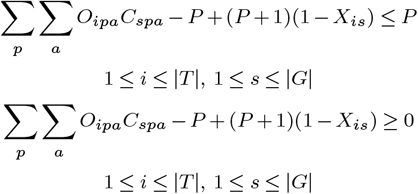

Finally, we can impose the following constraints to solve *t*_*i*_ for all values of *i* to ensure that it is covered by at least one degenerate template in *G*:

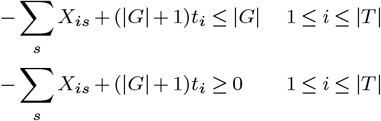

### Total library size constraint

We note that the calculation of total produced library size, |span_DNA_(*G*)|, requires multiplication of the total possible number of incorporated residues at each position. Because multiplication of variables is a non-linear operation, we instead calculate the log of the span of *G* and introduce the following constraint against the technology-imposed diversity limit *M* to ensure that log(|span_DNA_(*G*)|) ≤ log(*M*) and, therefore | span_DNA_(*G*)| ≤ *M* in the case of a single sublibrary, *s* = 1:

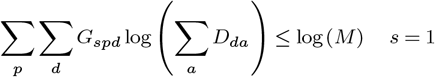

### MC-DMTD: Multiple degenerate templates extension

To extend the ILP for use with multiple degenerate templates, we must adjust the calculation of the constraint on the size of the total produced library. Because this calculation requires a sum of products, it is not trivial to devise a linear solution. Instead, we approximate the solution by binning the log of the sublibrary size produced by each degenerate template into discrete bins ranging from size 1 to size log(*M*). We then approximate the size of each sublibrary as the exponentiated value of the upper bound of the bin, thus ensuring that the calculated approximate sublibrary size is always greater than or equal to the true sublibrary size across all sublibraries. Therefore, the constraint that the total library size must be less than or equal to the user-defined limit will always hold for a valid solution of the ILP.

To calculate the appropriate bin for the size of the sublibrary produced by each degenerate template, we define two vectors *U* and *L* such that *U*_*n*_ and *L*_*n*_ define the upper and lower bound of bin *n*, respectively. We then let *Q*_*s*_ denote the log size of the sublibrary span_DNA_(*G*_*s*_) calculated as *Q*_*s*_ = Σ_*p*_ Σ_*d*_ *G*_*spd*_ log (Σ_*a*_ *D*_*da*_). We then introduce a binary-valued variable *B*_*sn*_ to indicate whether the size of sublibrary s falls into the range of bin *n* such that *L*_*n*_ ≤ log(|span_DNA_(*G*_*s*_)|) ≤ *U*_*n*_. Upon *B*_*sn*_, we place the following constraints:

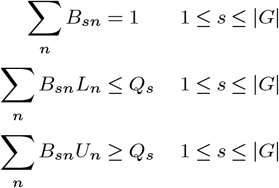

Our upper bound on total library size when combining multiple sublibraries is therefore calculated as:

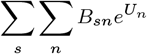

The following constraint can be introduced to ensure that the maximum possible library size when combining multiple sublibraries does not exceed the user-defined limit:

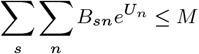

### Degenerate codon table

To reduce unnecessary search over equivalent solutions, we formulate the degenerate codon table *D* to minimize the degeneracy of the constructed DNA library while maintaining access to all 20 amino acid residues, stop codons, and gaps. Though 3,376 possible degenerate codons exist (15^3^ plus the empty set to account for a gap), many of these codons are redundant in the amino acids they encode. We therefore employ a non-redundant codon table that groups all codons covering the same set of amino acids. We select and return from this set the subset of codons that display the least degeneracy in nucleic acid space. By removing redundant codons from the codon table, we reduce the number of possible codon selections from 3,376 to 841.

### Implementation

DeCoDe was implemented in Python using the CVXPY package (8). All results presented here employed the Gurobi solver (12), which provides a free licence to academic users. CVXPY also provides interfaces to most common alternative ILP solvers including free and opensource options. All results presented here were run on a server with two Intel Xeon CPU E5-2630 v4 @ 2.20GHz CPUs and 256 GB of memory. Each run of the DeCoDe algorithm was allocated 12 hyper-threads for the Gurobi solver and multiple runs were conducted in parallel using GNU Parallel (41). We provide a command line interface to DeCoDe at github.com/OrensteinLab/DeCoDe.

## Results

We sought to apply DeCoDe to a set of library optimization problems closely mimicking those required for a standard protein engineering project. Specifically, we chose the task of covering as many sequences as possible from the documented lineage of the green fluorescent protein originally extracted from the jellyfish *Aequorea victoria* (avGFP) (30). We selected this task for several reasons. First, avGFP and its laboratory-derived descendents have provided a critical suite of research tools and, consequently, are the subject of ongoing research to map the protein’s sequence onto its functional characteristics such as intensity and stability (35). Second, the exact lineage of the entire family of laboratory-derived avGFP variants is known, which makes the task of aligning all of the sequences and identifying variations from wild type avGFP a trivial task (18). Third, the high total number of variable sites and the length variation within the protein family causes this design problem to be particularly challenging or impossible for existing DC library design algorithms.

### MC-DSTD performance

First, we compared DeCoDe to an existing degenerate codon library design algorithm to benchmark its performance. As the most recently published method and the only open-source algorithm compatible with multiple degenerate codon usage per position, SwiftLib (14) represents a natural point of comparison.

The total set of avGFP-derived proteins spans several different lengths, the most common being length 239 amino acids with all other proteins except two being of length 238. The two exclusive proteins include large, unique insertions and were excluded from our benchmarks since each would need to be ordered as individual, non-degenerate constructs in any degenerate codon library. While DeCoDe permits libraries of varying length, SwiftLib requires all target proteins to be the same length. To allow for a direct comparison between the two methods, we subset the GFP lineage to only proteins of length 239 amino acids.

We performed library optimization for this reduced target protein set with both DeCoDe and SwiftLib. As both algorithms are capable of increasing performance by employing multiple degenerate DNA templates, we optimized for a range of library size limits using either 1 or 2 sublibraries, where each sublibrary represents an independently synthesized DNA construct. We then measured the quality of the each algorithm’s generated library by the number of full length sequences out of the target set of 94 proteins covered by that DC library.

For library designs composed of only a single sublibrary, we find that DeCoDe consistently outperforms SwiftLib in the total number of target sequences covered for a given library size limit (Figure 2A). Across all limits for a single sublibrary, DeCoDe offers an improvement of between 58% and 250%. In both cases, the returned library sizes approach, but do not exceed, the user-specified limit (Figure S1). The superior performance of DeCoDe is a direct result of the objective function implicitly accounting for linked variation between multiple positions, whereas SwiftLib minimizes the failure to include a desired amino acid for each position independently. Still, DeCoDe’s runtime and memory usage are feasible by today’s standard: only around 100 and 1000 seconds runtime and around 1 and 2.5GB for 1 and 2 sublibrary designs, respectively.

**Fig 2.**
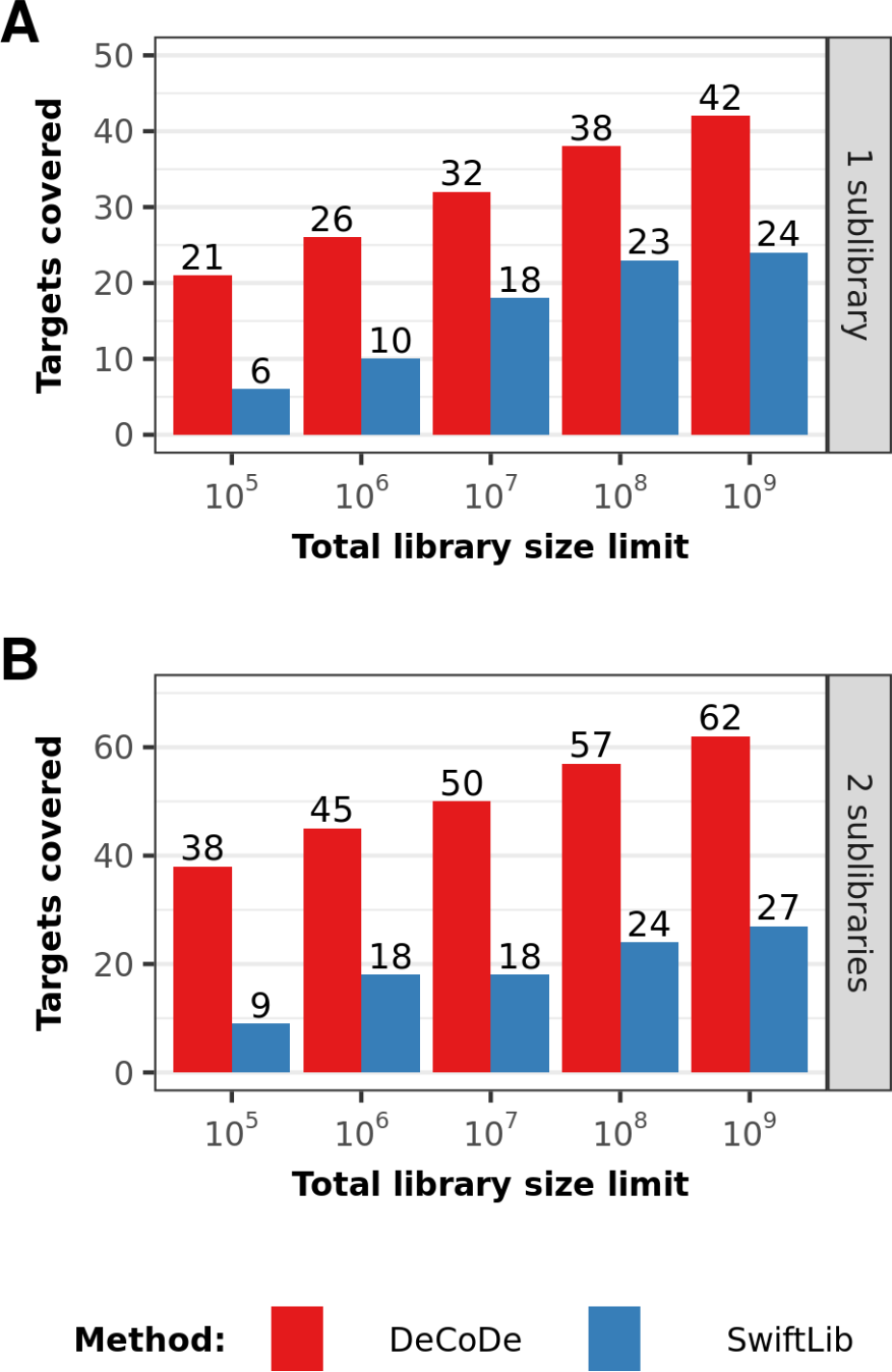
Comparison of target coverage by DeCoDe and SwiftLib on the target sequence set of 94 avGFP-descended target proteins of length 239 amino acids, where 82 positions are variable between proteins. Optimized libraries for each method are shown comprising (A) 1 sublibrary; and (B) 2 sublibraries.

By accounting for covariation between sites, DeCoDe can also more efficiently include multi-site patterns in the design of the DC libraries. In the case where DeCoDe is allowed to include a second sublibrary, and therefore better account for covariation, its performance boost over SwiftLib is even more dramatic (Figure 2B). For all diversity limits, DeCoDe outputs a library that covers more than twice the number of proteins coverd by SwiftLib’s library. However, because this problem is NP-hard, both the runtime (Figure S2) and memory requirements (Figure S3) of DeCoDe tend to be much higher than those of SwiftLib. Because SwiftLib does not attempt to optimize over the problem of covariation between multiple sites, the algorithm can find an optimal solution for its library quality metric in polynomial time.

### MC-DMTD performance

In addition to the 94 proteins of length 239 amino acids explored in the above optimization task, the avGFP family includes 37 additional proteins of length 238 amino acids. Because DeCoDe can employ multiple sublibraries and the gap codon, it is, to our knowledge, the first algorithm able to perform total library optimization for libraries composed of mixed length protein targets. Here, we use DeCoDe to optimize a library targeting the set of 131 avGFP-derived proteins of both 238 and 239 amino acid lengths (Figure 3).

**Fig 3.**
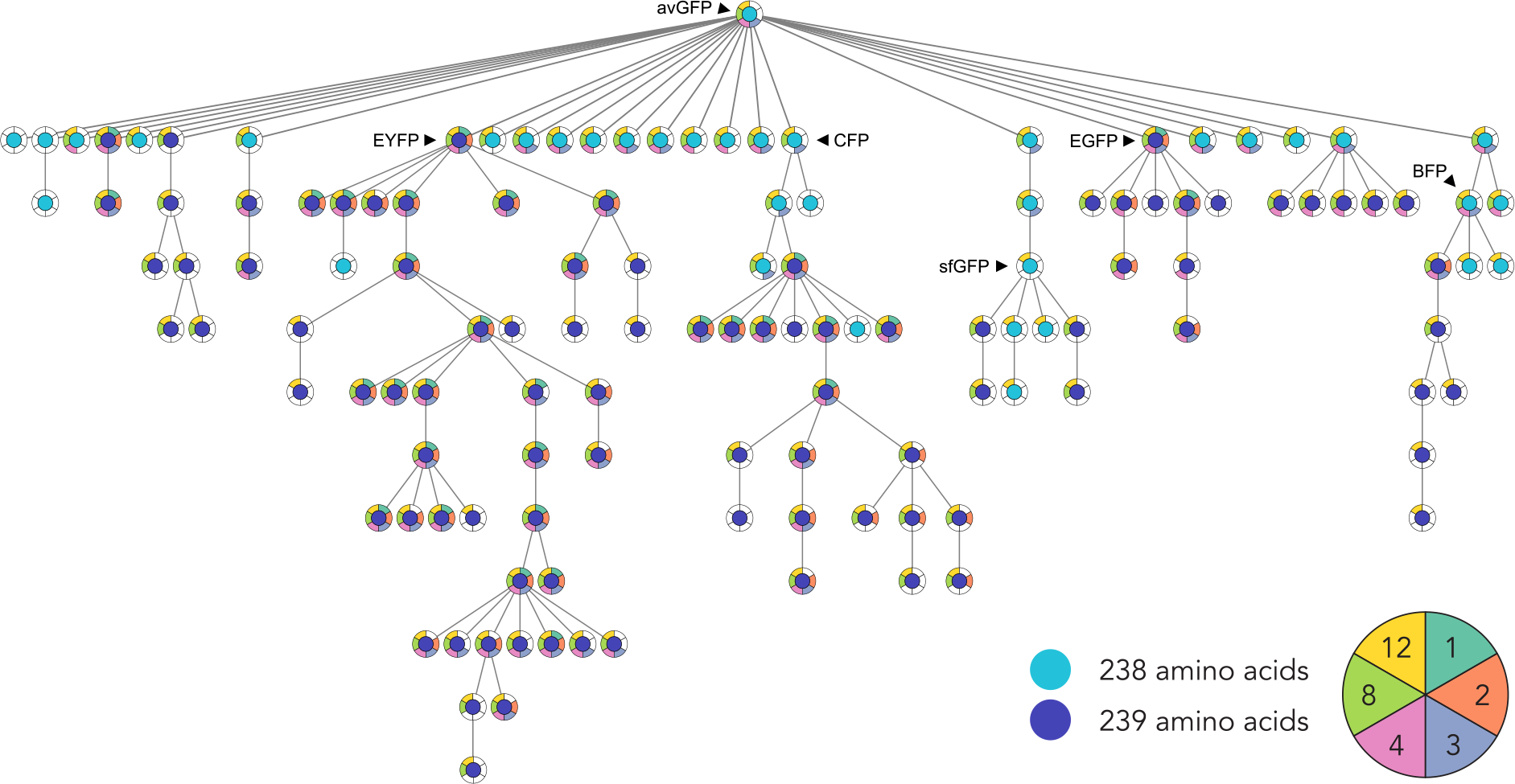
A phylogeny of 131 avGFP-descended proteins is shown. 94 and 37 proteins of length 239 and 238 amino acids are shown by circles colored dark and light blue, respectively. The colored segments behind each circle denote coverage by libraries comprising the number of sublibraries shown in the key in the lower right of the figure. Important avGFP descendants exhibiting spectral variation (EYFP, CFP, BFP), intensity enhancement (EGFP), and rapid maturation (sfGFP) are labelled on the tree.

For this task, we selected a total library size limit of 10^7^ unique DNA species. This limit is consistent with yeast-based assays (*e.g.* yeast display) frequently used in conjunction with fluorescence-activated cell sorting (FACS) to enrich for fluorescent proteins with desirable properties (39). Because the avGFP family is relatively diverse, we explored different sublibrary counts to maximize coverage under the 10^7^ diversity limit. Specifically, we optimized libraries comprising 1, 2, 3, 4, 8, and 12 sublibraries. Each sub-library represents an additional monetary and labor cost that is, in most instances, offset by the improved coverage that the sublibraries achieve under the total diversity limit. Because each additional sublibrary further complicates the optimization procedure, libraries comprised of a high number of sublibraries will likely require a prohibitively long runtime to reach a guaranteed optimal solution. To address this issue, we limited the total runtime to a maximum of 48 hours and selected the solution reached by the ILP solver under that time limit.

The inclusion of additional sublibraries dramatically improved overall coverage of the lineage tree (Figure S4) while keeping the total library diversity under the 10^7^ limit (Figure S5). When only a single sublibrary is used, DeCoDe outputs an optimal library. However, this library only covers 32 of the 131 target sequences and all of the covered sequences are of length 239. When 2 additional sublibraries are added (for a total of 3), the library produced by DeCoDe covers 66 proteins, or just over half of all sequences in the lineage. This 3-member library begins to cover sequences of length 238 with diverse functions, including CFP and BFP. With 12 sublibraries, DeCoDe covers 121 of the 131 total target sequences while generating only 7,640,832 total DNA sequences, which is well within the range of modern DNA synthesis techniques. With this library, nearly all of the significant functional variants are covered. As expected, there is a positive correlation between the number of sublibraries included and the runtime (Figure S6) and maximum memory usage (Figure S7) for each optimization problem.

By further exploring the covariation structure of the avGFP lineage tree and comparing it with the patterns of coverage obtained when adding additional sublibraries, several interesting patterns emerge (Figure 3). In the case of a single sublibrary, only proteins of length 239 amino acids are covered. This result is expected as there are more sequences of length 239 (94 sequences) than those of length 238 (37 sequences). Because the gap codon covers the gap and only the gap, an additional sublibrary must be ordered for each additional desired protein length.

Second, entire branches of the lineage tend to be covered together. Because sequences lower in the tree share a common ancestor, derived sequences are likely clustered together within the coverage space of a single or a small subset of sublibraries. However, not all derived sequences are covered when the algorithm begins to include a branch of the tree. DeCoDe may determine that covering a single specific sequence within a branch may incur a penalty toward the total diversity limit that is too great, and that sequence may be excluded. In some cases, such as that of some terminal branches of the CFP-derived lineage, certain protein sequences are covered with a lower number of sublibraries (e.g. 2) and not covered when more are added (e.g. 3, 4). This pattern emerges when it becomes optimal for the algorithm to allocate sublibraries to more tightly clustered groups of sequences when additional sublibraries are added.

When employing a smaller number of sublibraries (i.e. 1, 2, 3), DeCoDe returns a solution that samples shallowly from many branches within the tree. Therefore, a greater diversity in potential functional outcomes is explored at the expense of a limit on the total number of target sequences covered. In contrast, with a large number of sublibraries (i.e. 4, 8, 12), the algorithm covers large branches more completely due to the smaller total diversity for each individual sublibrary. The major tradeoffs when employing a greater number of sublibraries are the increased computational costs to optimize the library and the increased expense of ordering and processing multiple synthesized DNA constructs. Therefore, the user must ultimately decide on an optimal point between target coverage and computational and experimental cost.

While we were able to obtain guaranteed optimal solutions for the case of 1 and 2 sublibraries, we were unable to obtain optimality guarantees for the libraries comprising 3, 4, 8, and 12 sublibraries under the 48 hour time limit. However, solutions tend to rapidly improve on the objective function and yield diminishing returns with increased runtimes (Figure S8). The ILP solver computes an upper bound for the value of the optimal solution for each point during the run, which can inform the user how far the current library is from a theoretical optimum. Given these findings, we suggest that DeCoDe is unlikely to significantly improve on the presented results with increased time and that reaching a guaranteed optimal solution may require a dramatic increase in runtime.

## Discussion

The synthesis of large-scale protein-coding DNA libraries is often a necessary first step for high-throughput protein screening assays. This synthesis step often incurs a high cost, even for libraries of closely related genes. Many research groups have focused significant efforts on reducing this prohibitive cost through innovations in software (14, 24, 27), hardware (20, 26), or chemistry (29). Among these techniques, DC libraries stand out as a particularly attractive method, as they can cover large swaths of sequence space without a proportional rise in synthesis cost. However, existing DC library design solutions lack the ability to account for either linked variation, multiple protein lengths, or both. DeCoDe was designed with careful consideration of both linked and length variation, making it a particularly useful solution for a variety of DC library design tasks.

While these design choices enable DeCoDe to tackle previously untenable challenges, DeCoDe’s direct solution for the NP-hard problem of linked variation can require significant computational resources to find optimal libraries. We implemented two features to overcome these limits. The first is the use of a non-redundant codon table in the ILP formulation, which reduces the number of constraints and variables to consider. The second is a user-defined limit on the runtime of the ILP solver. While the ILP may not find an optimal solution with limited runtime, it will output a feasible solution that may be very close to the optimum.

DeCoDe is best suited for optimization tasks with high sequence similarity across targets and significant covariation between amino acid positions for high-throughput screening. For example, in the task of optimizing a library to cover the avGFP-derived family presented here, only 82 of the 239 possible positions varied. In the case of highly diverse libraries where the majority of positions are variable, DC libraries may provide low target coverage while simultaneously producing a high number of off-target sequences. In target libraries with mostly independent positions, algorithms which assume independence, such as SwiftLib (14), may be more appropriate.

Covariation between amino acid positions underlies the evolutionary structure of protein families. Naturally evolved proteins rely on evolutionarily conserved, interconnected networks of residue interactions to carry out their functions (13, 23, 38). These networks can often be disrupted by even a single amino acid change if that change is non-conservative in a necessary physical property (11). Several research groups have exploited the preservation of these networks over time to reconstruct ancestral protein lineages and better understand the link between protein sequence structure and function (22, 36, 43). Due to the highly correlated, chemically conserved sequence patterns present in these reconstructed lineages, DeCoDe offers an attractive solution to the problem of synthesizing the complete protein family for functional testing simultaneously.

Expansions or reductions in protein domain lengths represent another type of variation with significant implications for protein function (6). As an example, size differences in the complementarity-determining regions (CDRs) of immunoglobulin proteins can differentiate success and failure of antigen binding (42). Because DeCoDe can simultaneously optimize constructs of various lengths under a single library size constraint, it is ideal for generating DC libraries to screen immunoglobulin proteins for specific, desired binding properties.

Given a set of proteins with unknown functionality, it is highly likely that a final library designed by DeCoDe will include variants with novel, and perhaps useful, functions. Libraries produced by DeCoDe will be most useful when the screening method employed by the user is tolerant to offtarget proteins, as many functional variants may reside in the space of the “off-target” sequences not present in the target set. DeCoDe-generated libraries stand to make a significant impact in the fields of directed evolution and ancestral reconstruction as they have the capacity to more efficiently screen sequence space for functional variants. Moreover, the cost reduction achieved by the use of DeCoDe’s more efficient DC libraries enables greater experimental throughput and a more rapid pace of functional protein discovery.

## Acknowledgements

TCS acknowledges travel support from The Prof. Rahamimoff Travel Grant Program of the United States-Israel Binational Science Foundation (BSF). TCS acknowledges the support of an NSF Graduate Research Fellowship. PMF is a Chan Zuckerberg Biohub Investigator and acknowledges the support of an Alfred P. Sloan Foundation Fellowship. This preprint is formatted using a LATEX class by Ricardo Henriques.

## References

1. Agresti, J. J. et al. (2010). Ultrahigh-throughput screening in drop-based microfluidics for directed evolution. Proc. Natl. Acad. Sci. U.S.A., 107, 4004–4009.

2. Arkin, A. P. and Youvan, D. C. (1992). Optimizing nucleotide mixtures to encode specific subsets of amino acids for semi-random mutagenesis. Nat. Biotechnol., 10, 297–300.

3. Barbas, C. F. et al. (1991). Assembly of combinatorial antibody libraries on phage surfaces: the gene III site. Proc. Natl. Acad. Sci. U.S.A., 88(18), 7978–7982.

4. Beaucage, S. L. and Caruthers, M. H. (1981). Deoxynucleoside phosphoramidites - a new class of key intermediates for deoxypolynucleotide synthesis. Tetrahedron Lett., 22, 1859–1862.

5. Boder, E. T. and Wittrup, K. D. (1997). Yeast surface display for screening combinatorial polypeptide libraries. Nat. Biotechnol., 15, 553–557.

6. Brocchieri, L. and Karlin, S. (2005). Protein length in eukaryotic and prokaryotic proteomes. Nucleic Acids Res., 33, 3390–3400.

7. Cadet, F. et al. (2018). A machine learning approach for reliable prediction of amino acid interactions and its application in the directed evolution of enantioselective enzymes. Sci. Rep., 8, 16757.

8. Diamond, S. and Boyd, S. (2016). CVXPY: A python-embedded modeling language for convex optimization. Journal of Machine Learning Research, 17, 221–264.

9. Eisenmesser, E. Z. et al. (2005). Intrinsic dynamics of an enzyme underlies catalysis. Nature, 438, 117–121.

10. Freudl, R. et al. (1986). Cell surface exposure of the outer membrane protein OmpA of Escherichia coli K-12. J. Mol. Biol., 188, 491–494.

11. Goldberg, A. L. and Wittes, R. E. (1966). Genetic code: aspects of organization. Science, 153(3734), 420–424.

12. Gurobi Optimization, L. (2018). Gurobi optimizer reference manual.

13. Halabi, N. et al. (2009). Protein sectors: evolutionary units of three-dimensional structure. Cell, 138(4), 774–786.

14. Jacobs, T. M. et al. (2014). SwiftLib: rapid degenerate-codon-library optimization through dynamic programming. Nucleic Acids Res., 43(5), e34–e34.

15. Jiang, L. et al. (2008). De novo computational design of retro-aldol enzymes. Science, 319, 1387–1391.

16. Kuhlman, B. et al. (2003). Design of a novel globular protein fold with atomic-level accuracy. Science, 302, 1364–1368.

17. LaBean, T. H. and Kauffman, S. A. (1993). Design of synthetic gene libraries encoding random sequence proteins with desired ensemble characteristics. Protein Sci., 2, 1249–1254.

18. Lambert, T. J. (2019). FPbase: a community-editable fluorescent protein database. Nat. Methods, 16, 277–278.

19. Leaver-Fay, A. et al. (2011). ROSETTA3: an object-oriented software suite for the simulation and design of macromolecules. In Methods in Enzymology, volume 487, pages 545–574. Elsevier.

20. LeProust, E. M. et al. (2010). Synthesis of high-quality libraries of long (150mer) oligonucleotides by a novel depurination controlled process. Nucleic Acids Res., 38(8), 2522–2540.

21. Lewis, S. M. et al. (2014). Generation of bispecific IgG antibodies by structure-based design of an orthogonal Fab interface. Nat. Biotechnol., 32, 191–198.

22. Lim, S. A. et al. (2016). Evolutionary trend toward kinetic stability in the folding trajectory of RNases H. Proc. Natl. Acad. Sci. U.S.A., 113, 13045–13050.

23. Lockless, S. W. and Ranganathan, R. (1999). Evolutionarily conserved pathways of energetic connectivity in protein families. Science, 286, 295–299.

24. Mena, M. A. and Daugherty, P. S. (2005). Automated design of degenerate codon libraries. Protein Engineering, Design & Selection, 18(12), 559–561.

25. Motlagh, H. N. et al. (2014). The ensemble nature of allostery. Nature, 508, 331–339.

26. Oling, D. et al. (2018). Large scale synthetic site saturation GPCR libraries reveal novel mutations that alter glucose signaling. ACS Synth. Biol., 7(9), 2317–2321.

27. Parker, A. S. et al. (2011). Optimization of combinatorial mutagenesis. J. Comput. Biol., 18(11), 1743–1756.

28. Pierce, N. A. and Winfree, E. (2002). Protein design is NP-hard. Protein Eng., 15, 779–782.

29. Plesa, C. et al. (2018). Multiplexed gene synthesis in emulsions for exploring protein functional landscapes. Science, 359, 343–347.

30. Prasher, D. C. et al. (1992). Primary structure of the Aequorea victoria green-fluorescent protein. Gene, 111, 229–233.

31. Roberts, R. W. and Szostak, J. W. (1997). RNA-peptide fusions for the in vitro selection of peptides and proteins. Proc. Natl. Acad. Sci. U.S.A., 94, 12297–12302.

32. Rockberg, J. et al. (2008). Epitope mapping of antibodies using bacterial surface display. Nat. Methods, 5, 1039–1045.

33. Romero, P. A. et al. (2015). Dissecting enzyme function with microfluidic-based deep mutational scanning. Proc. Natl. Acad. Sci. U.S.A., 112(23), 7159–7164.

34. Saito, Y. et al. (2018). Machine-learning-guided mutagenesis for directed evolution of fluorescent proteins. ACS Synth. Biol., 7(9), 2014–2022.

35. Sarkisyan, K. S. et al. (2016). Local fitness landscape of the green fluorescent protein. Nature, 533, 397–401.

36. Shi, Y. and Yokoyama, S. (2003). Molecular analysis of the evolutionary significance of ultraviolet vision in vertebrates. Proc. Natl. Acad. Sci. U.S.A., 100, 8308–8313.

37. Smith, G. (1985). Filamentous fusion phage: novel expression vectors that display cloned antigens on the virion surface. Science, 228, 1315–1317.

38. Socolich, M. et al. (2005). Evolutionary information for specifying a protein fold. Nature, 437, 512–518.

39. Swers, J. S. et al. (2004). Shuffled antibody libraries created by in vivo homologous recombination and yeast surface display. Nucleic Acids Res., 32, e36.

40. Tabuchi, I. et al. (2001). An in vitro DNA virus for in vitro protein evolution. FEBS Lett., 508(3), 309–312.

41. Tange, O. (2018). GNU Parallel 2018. Ole Tange, first edition.

42. Teplyakov, A. and Gilliland, G. L. (2014). Canonical structures of short CDR-L3 in antibodies. Proteins, 82, 1668–1673.

43. Thornton, J. W. et al. (2003). Resurrecting the ancestral steroid receptor: ancient origin of estrogen signaling. Science, 301, 1714–1717.

44. Wolf, E. and Kim, P. S. (1999). Combinatorial codons: a computer program to approximate amino acid probabilities with biased nucleotide usage. Protein Sci., 8, 680–688.

45. Wu, Z. et al. (2019). Machine learning-assisted directed protein evolution with combinatorial libraries. Proc. Natl. Acad. Sci. U.S.A., page 201901979.

