## Supplementary Information for "DeCoDe: degenerate codon design for complete protein-coding DNA libraries"

### Contents

**1 ILP Formulation Proofs** **3**

**2 Complete ILP formulations** **8**

2.1 Single template . . . . . . . . 8

2.2 Multiple templates . . . . . . . . 9

**3 Supplementary Figures** **10**

### 1 ILP Formulation Proofs

**Claim 1.** *$X_{is} = 1 \iff T_i \in span_{AA}(g_s)$ by the following constraints:*

$$\sum_p \sum_a O_{ipa} C_{spa} - P + (P+1)(1 - X_{is}) \leq P \quad 1 \leq i \leq |T|,\ 1 \leq s \leq |G|$$
$$\sum_p \sum_a O_{ipa} C_{spa} - P + (P+1)(1 - X_{is}) \geq 0 \quad 1 \leq i \leq |T|,\ 1 \leq s \leq |G|$$

*Proof.* We consider two scenarios. In one $0 \leq \sum_p \sum_a O_{ipa} C_{spa} \leq P - 1$. Then, $T_i$ is not covered by $g_s$, thus $X_{is}$ should be 0. Indeed, only $X_{is} = 0$ satisfies the two constraints. In the second scenario $\sum_p \sum_a O_{ipa} C_{spa} = P$, thus $X_{is}$ should be 1. Indeed, only $X_{is} = 1$ satisfies the two constraints. □

**Claim 2.** *$t_i = 1 \iff \exists g_s \in G, T_i \in span_{AA}(g_s)$ by the following constraints:*

$$-\sum_s X_{is} + (|G|+1)t_i \leq |G| \quad 1 \leq i \leq |T|$$
$$-\sum_s X_{is} + (|G|+1)t_i \geq 0 \qquad 1 \leq i \leq |T|$$

*Proof.* Consider the case that $T_i$ is not covered. Then, all $X_{is}$ are zero. Then, only $t_i = 0$ can satisfy the two constraints, as $0 \leq |G|$ and $0 \geq 0$. Consider the other case, i.e. $T_i$ is covered by at least one degenerate template, so $-|G| \leq -\sum_s X_{is} \leq -1$. In this case, only $t_i = 1$ can satisfy both constraints. □

**Lemma 1.** *For vector $x$ and constant matrix $A$,*
$(x_i \in \{0,1\} \ \forall i) \wedge (\sum_i x_i = 1) \wedge (\sum_j A_{ij} > 0 \ \forall i) \longrightarrow \log\left(\sum_i x_i \sum_j A_{ij}\right) = \sum_i x_i \log\left(\sum_j A_{ij}\right)$.

*Proof.* Let $\hat{\imath}$ indicate the index for which $x_{\hat{\imath}} = 1$:

$$\begin{aligned}
\sum_i x_i \log\left(\sum_j A_{ij}\right) &= \log\left(\sum_i x_i \sum_j A_{ij}\right) \\
\sum_{i \neq \hat{\imath}} x_i \log\left(\sum_j A_{ij}\right) + x_{\hat{\imath}} \log\left(\sum_j A_{\hat{\imath}j}\right) &= \log\left(\sum_{i \neq \hat{\imath}} x_i \sum_j A_{ij} + x_{\hat{\imath}} \sum_j A_{\hat{\imath}j}\right) \\
x_{\hat{\imath}} \log\left(\sum_j A_{\hat{\imath}j}\right) &= \log\left(x_{\hat{\imath}} \sum_j A_{\hat{\imath}j}\right) \qquad (1) \\
1 \log\left(\sum_j A_{\hat{\imath}j}\right) &= \log(1) + \log\left(\sum_j A_{\hat{\imath}j}\right) \\
\log\left(\sum_j A_{\hat{\imath}j}\right) &= \log\left(\sum_j A_{\hat{\imath}j}\right)
\end{aligned}$$

□

**Claim 3.** *$|\text{span}_{DNA}(G)| \leq M$ in the case of a single degenerate template, i.e. $s = 1$, by the following constraint:*

$$\sum_p \sum_d G_{spd} \log \left( \sum_a D_{da} \right) \leq \log(M) \quad s = 1$$

*Proof.* In the case where $|G| = 1$ and therefore $s = 1$:

$$\begin{aligned}
\text{span}_{\text{DNA}}(G) &= \prod_p \sum_d G_{spd} \sum_a D_{da} && \leq M \\
\log(\text{span}_{\text{DNA}}(G)) &= \sum_p \log \left( \sum_d G_{spd} \sum_a D_{da} \right) && \leq \log(M) \\
\log(\text{span}_{\text{DNA}}(G)) &= \sum_p \sum_d G_{spd} \log \left( \sum_a D_{da} \right) && \leq \log(M)
\end{aligned}$$

The transition from the second to the third line follows from Lemma 1.

□

To calculate the appropriate bin for the size of the sublibrary produced by each degenerate template, we define two vectors $U$ and $L$ such that $U_n$ and $L_n$ define the upper and lower bound of bin $n$, respectively. Let $Q_s$ denote the log size of the sublibrary $\text{span}_{\text{DNA}}(G_s)$ calculated as $Q_s = \sum_p \sum_d G_{spd} \log\left(\sum_a D_{da}\right)$. We then introduce a binary-valued variable $B$

**Claim 4.** $B_{sn} = 1 \iff L_n \leq \log(|\,\text{span}_{DNA}(G_s)|) \leq U_n$, *by the following constraints:*

$$\begin{aligned}
\sum_n B_{sn} &= 1 \qquad 1 \leq s \leq |G| \\
\sum_n B_{sn} L_n &\leq Q_s \quad 1 \leq s \leq |G| \\
\sum_n B_{sn} U_n &\geq Q_s \quad 1 \leq s \leq |G|
\end{aligned}$$

*Proof.* Suppose that the value $Q_s$ falls between the values of $L_j$ and $U_j$. By the first constraint, only one value of $B_s$ can be 1, with all others taking value 0. Then, by the second constraint, $B_{sn}$ must be 0 for any values of $n \geq j$. By the first constraint, $B_{sn}$ must be 0 for any values $n \leq j$. Therefore the only feasible assignment is $B_{sj} = 1$. □

We note that because bin boundaries are inclusive, there exist two valid solutions when the value of $Q_s$ is equivalent to a bin boundary. In this instance, we rely on the ILP solver to select the lower-valued bin when necessary to provide a more accurate estimate of total library size.

### 2 Complete ILP formulations

#### 2.1 Single template

$$
\begin{aligned}
\text{max} \quad & \sum_i t_i \\
\text{subject to} \quad & \sum_d G_{spd} = 1 && \forall s, p \\
& \sum_p \sum_a O_{ipa} C_{spa} - P + (P+1)(1 - X_{is}) \leq P && \forall i, s \\
& \sum_p \sum_a O_{ipa} C_{spa} - P + (P+1)(1 - X_{is}) \geq 0 && \forall i, s \\
& -\sum_s X_{is} + (|G|+1) t_i \leq |G| && \forall i \\
& -\sum_s X_{is} + (|G|+1) t_i \geq 0 && \forall i \\
& \sum_p \sum_d G_{spd} \log\left(\sum_a D_{da}\right) \leq \log(M) && s = 1
\end{aligned}
$$

##### 2.2 Multiple templates

$$
\begin{aligned}
\text{max} \quad & \sum_i t_i \\
\text{subject to} \quad & \sum_d G_{spd} = 1 && \forall s, p \\
& \sum_p \sum_a O_{ipa} C_{spa} - P + (P+1)(1 - X_{is}) \leq P && \forall i, s \\
& \sum_p \sum_a O_{ipa} C_{spa} - P + (P+1)(1 - X_{is}) \geq 0 && \forall i, s \\
& -\sum_s X_{is} + (|G|+1) t_i \leq |G| && \forall i \\
& -\sum_s X_{is} + (|G|+1) t_i \geq 0 && \forall i \\
& \sum_n B_{sn} = 1 && 1 \leq s \leq |G| \forall s \\
& \sum_n B_{sn} L_n \leq Q_s && 1 \leq s \forall s \\
& \sum_n B_{sn} U_n \geq Q_s && 1 \leq \forall s \\
& \sum_s \sum_n B_{sn} e^{U_n} <= M
\end{aligned}
$$

### 3 Supplementary Figures

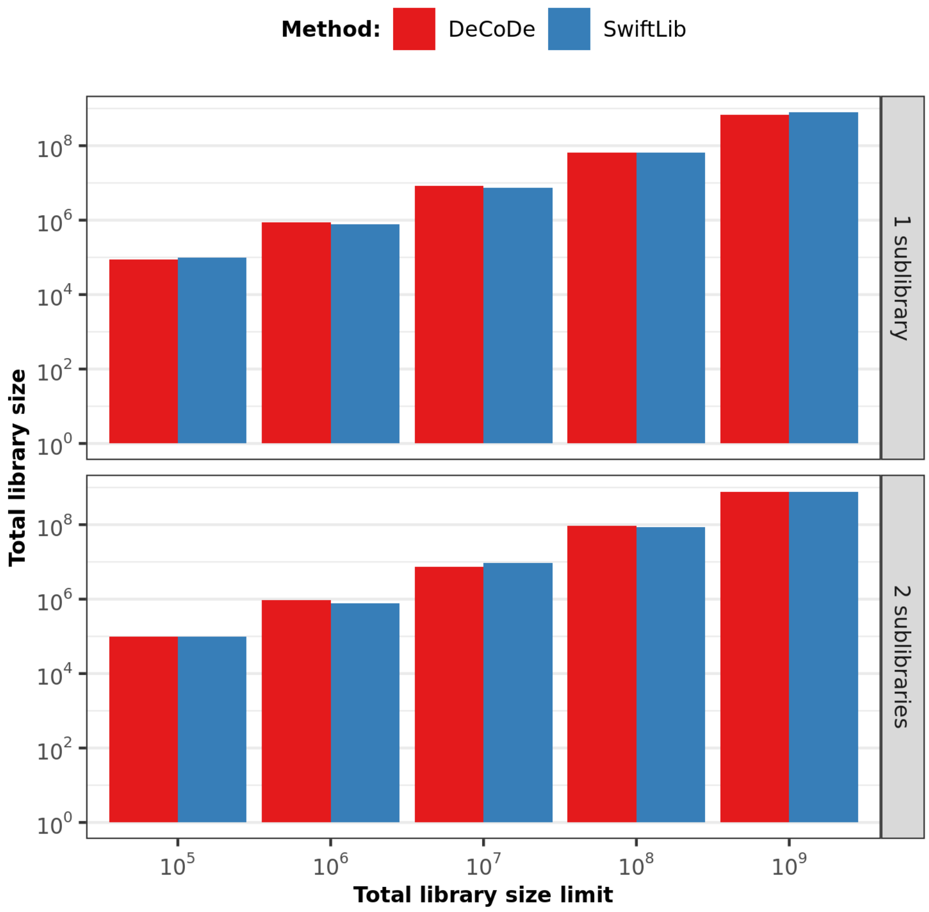


**Figure S1:** Total library size for both DeCoDe and Swiftlib for the task of maximizing coverage over the 94 avGFP-derived proteins of length 239 amino acids, where 82 positions vary between proteins. The limit for total library diversity is shown on the x-axis and the actual algorithm-utilized diversity is shown on the y-axis. In all cases, the actual diversity almost reaches the limit, but never exceeds it. Note both axes are on log scale.

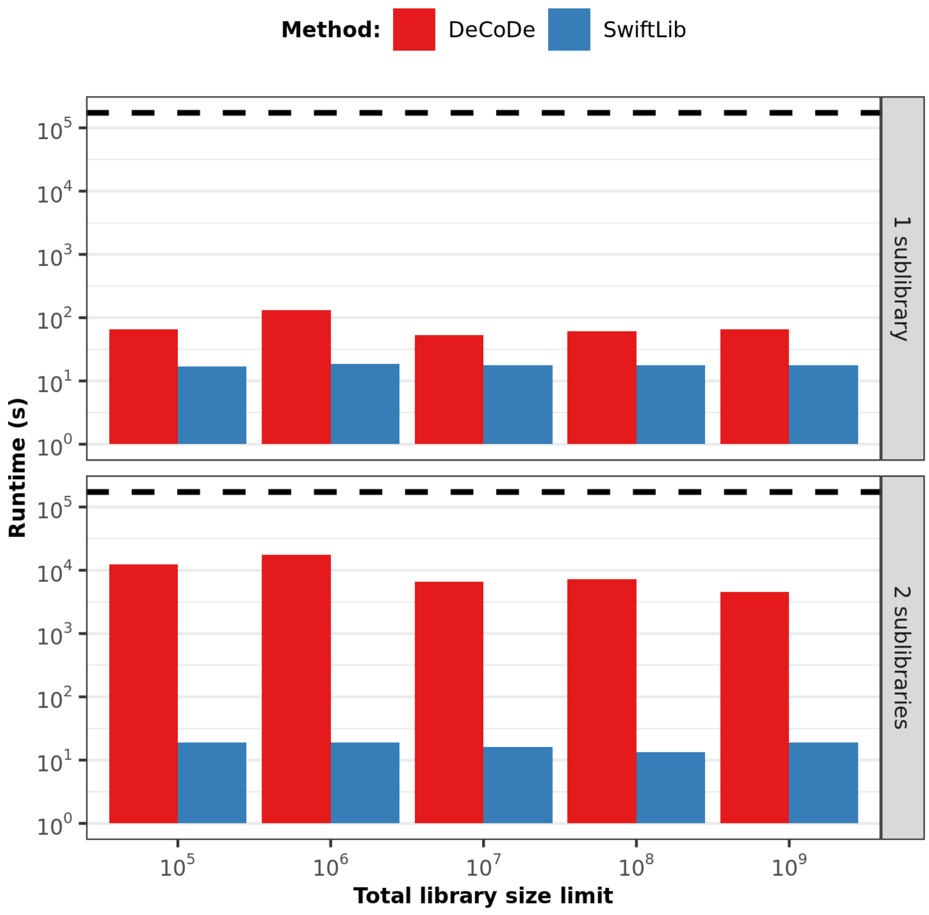


**Figure S2:** Total runtime for both DeCoDe and Swiftlib for the task of maximizing coverage over the 94 avGFP-derived proteins of length 239 amino acids, where 82 positions vary between proteins. The limit for total library diversity is shown on the x-axis and the runtime in seconds is shown on the y-axis. Note that both axes are on log scale. The total runtime of each run was limited to a maximum of 48 hours ($10^{5.24}$ seconds, as indicated by the black dashed line), though none of the runs encountered this time limit. The runtime for SwiftLib also includes the time necessary to start the Firefox web browser and load the server's webpage and is, therefore, an overestimate of total method-required runtime.

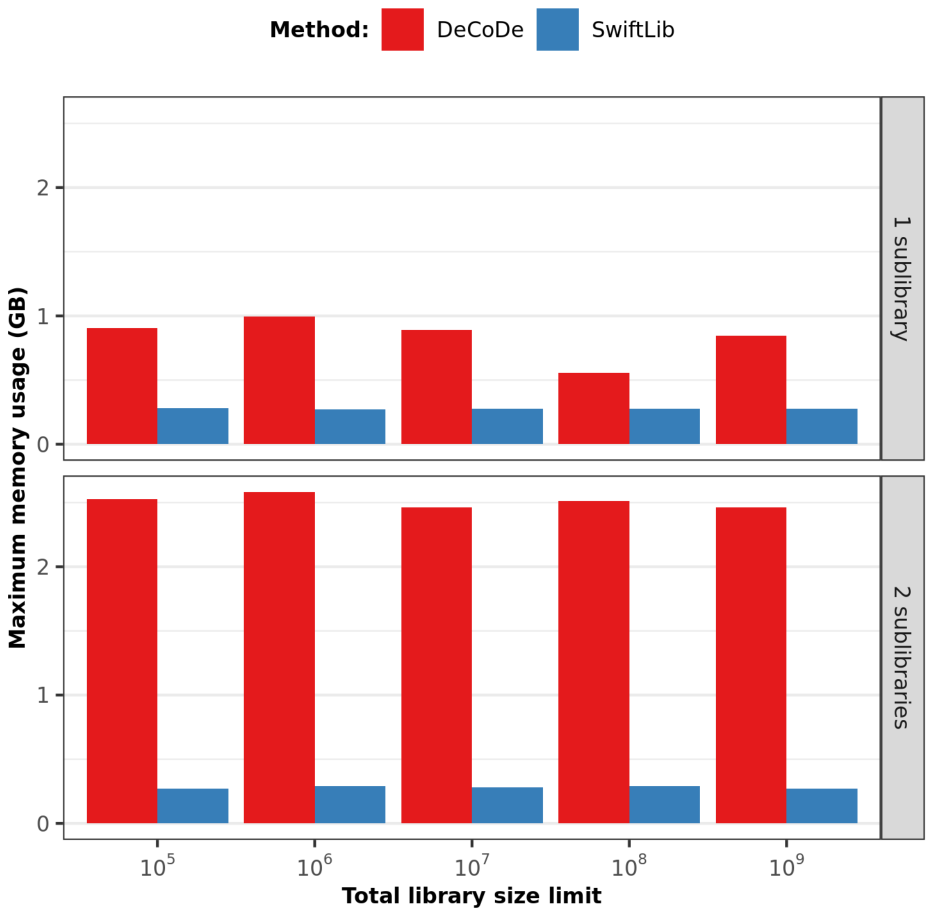


**Figure S3:** Maximum memory usage for both DeCoDe and Swiftlib for the task of maximizing coverage over the 94 avGFP-derived proteins of length 239 amino acids, where 82 positions vary between proteins. The limit for total library diversity is shown on the x-axis and the maximum memory usage in gigabytes is shown on the y-axis. Note that the x-axis is in log scale. The memory usage for SwiftLib also includes the memory necessary to start the Firefox web browser and load the server's webpage and, therefore, may overestimate total method-required memory.

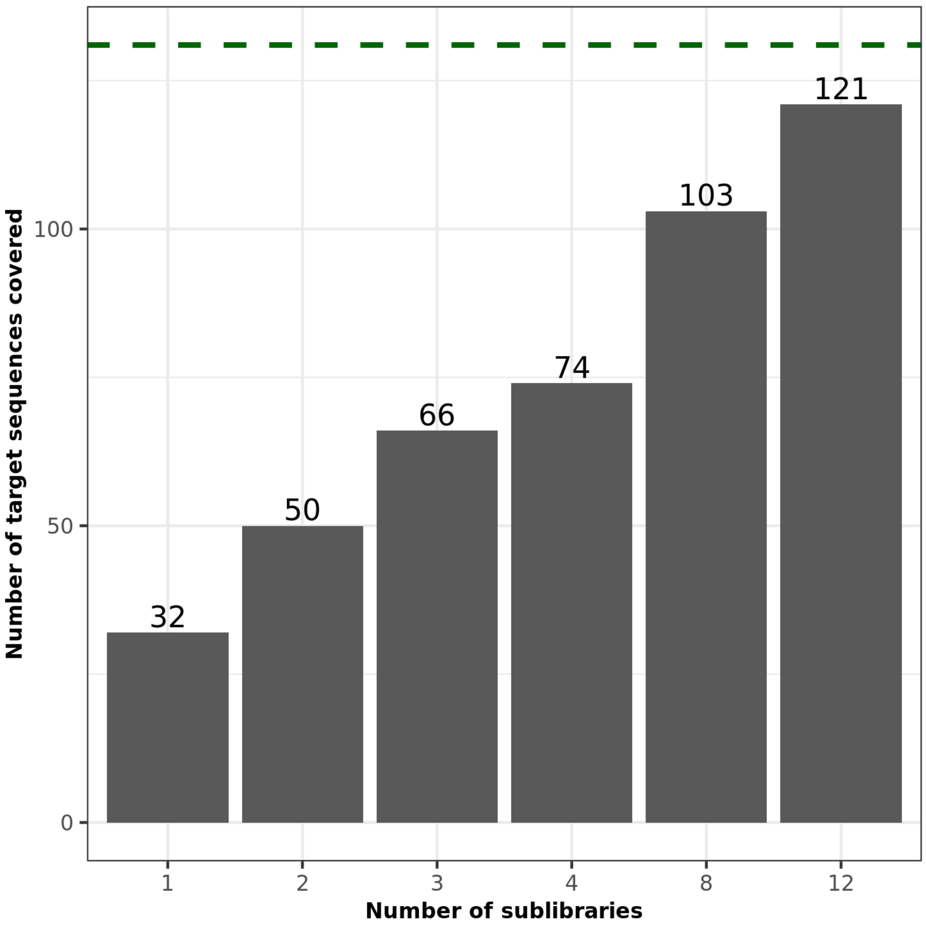


**Figure S4:** Total target library coverage produced by DeCoDe for the task of maximizing coverage over all 238 or 239 amino acid long avGFP-derived proteins under a total diversity limit of $10^7$ total possible DNA species. The number of degenerate templates is shown on the x-axis and the total number of covered target sequences is shown on the y-axis. The maximum possible number of covered sequences is 131 as indicated by the green dashed line.

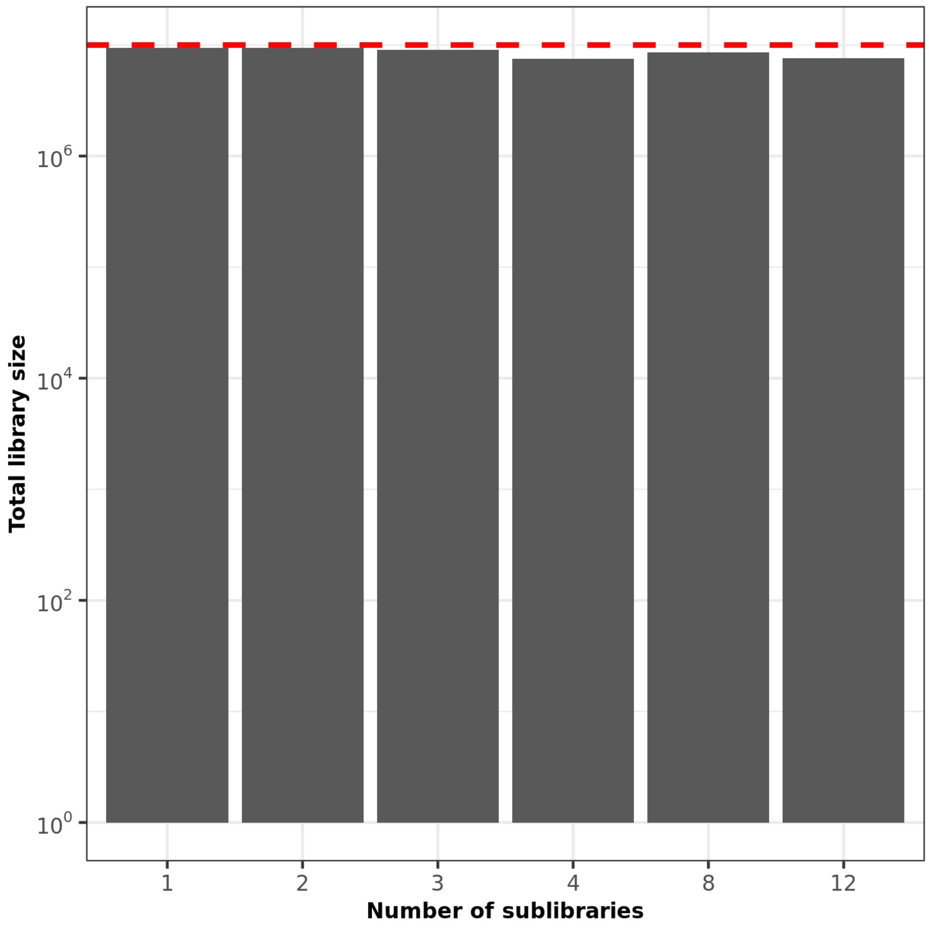


**Figure S5:** Total library size produced by DeCoDe for the task of maximizing coverage over all 238 or 239 amino acid long avGFP-derived proteins under a total diversity limit of $10^7$ total possible DNA species. The total number of constituent sublibraries is shown on the x-axis and the total number of covered target sequences is shown on the y-axis in log scale. In all cases, the actual diversity almost reaches the diversity limit of $10^7$ (indicated by the red dashed line), but never exceeds it.

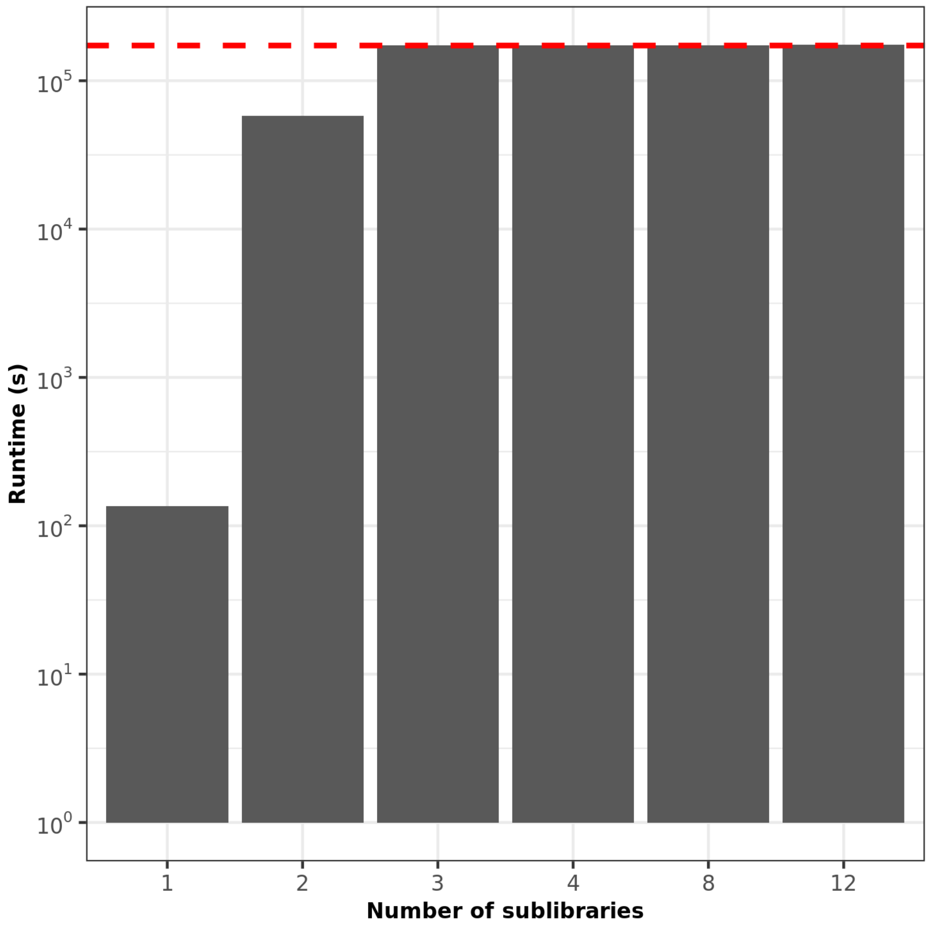


**Figure S6:** Total runtime of DeCoDe for the task of maximizing coverage over all 238 or 239 amino acid long avGFP-derived proteins under a total diversity limit of $10^7$ total possible DNA species. The total number of degenerate templates is shown on the x-axis and the runtime in seconds is shown on the y-axis in log scale. All ILP processes were terminated after a total runtime of 48 hours, corresponding to $10^{5.24}$ (indicated by the red dashed line) seconds even if they had not reached a guaranteed optimal solution.

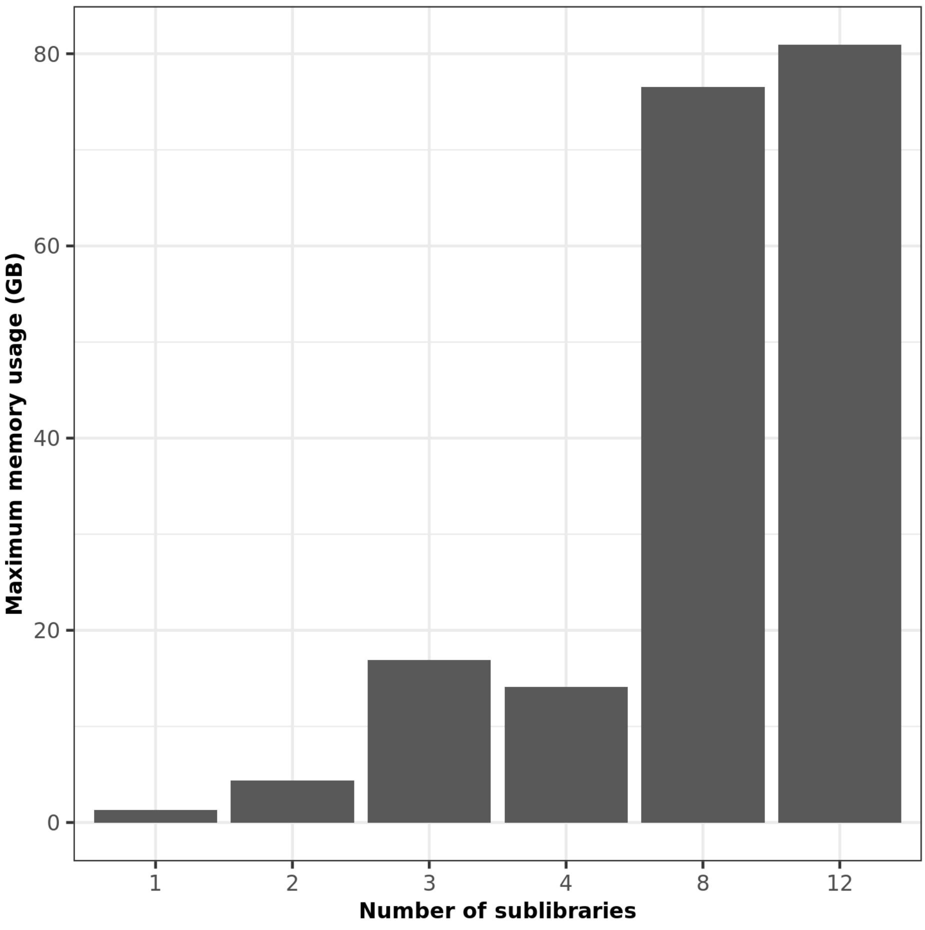


**Figure S7:** Maximum memory usage of DeCoDe for the task of maximizing coverage over all 238 or 239 amino acid long avGFP-derived proteins under a total diversity limit of $10^7$ total possible DNA species. The total number of degenerate templates is shown on the x-axis and the maximum memory usage in gigabytes is shown on the y-axis.

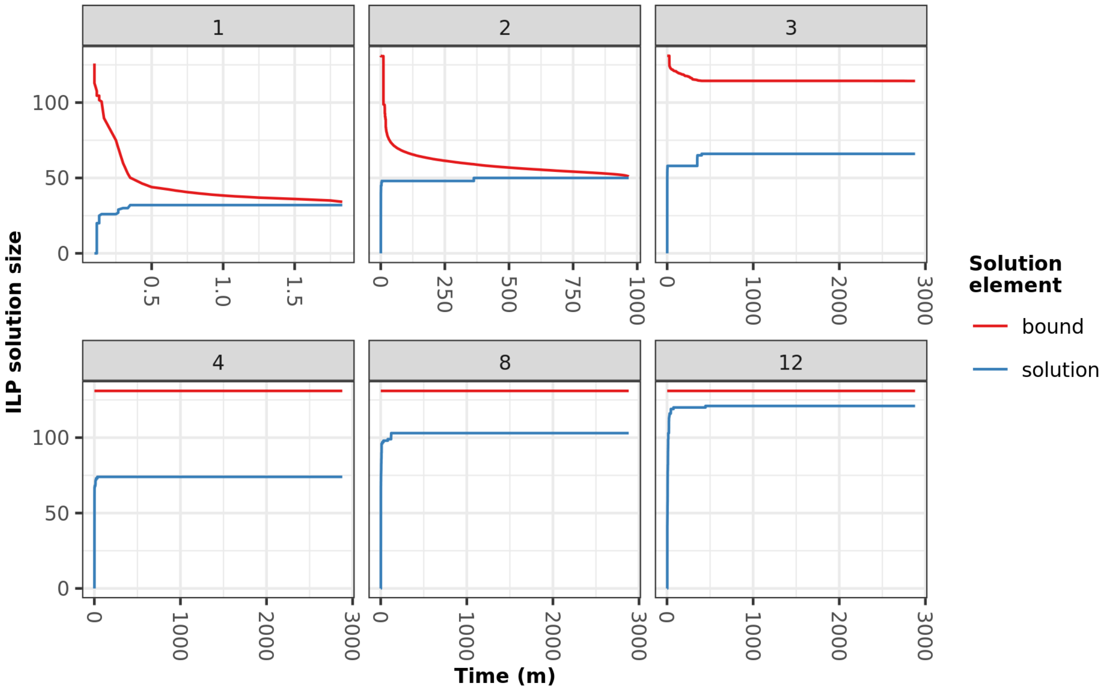


**Figure S8:** The gap between the current solution and the computed upper bound of an optimal solution is shown over time for each sublibrary count. The values of the coverage of the input library and the upper bound for optimality at each time shown by the blue and red lines, respectively. For the libraries consisting of one or two degenerate templates, the ILP solver eventually finds the optimal solution, resulting in no gap between these two values and termination of the solver before the 48hr time limit. However, for the remaining instances of multiple templates (three or more), the solver was unable to guarantee that the solution it reached is indeed optimal within the imposed time limit (48 hours).
